# Impact of galvanic vestibular stimulation electrode current density on brain current flow patterns: Does electrode size matter?

**DOI:** 10.1101/2022.08.17.504344

**Authors:** Dennis Q Troung, Alexander Guillen, Mujda Nooristani, Maxime Maheu, Francois Champoux, Abhishek Datta

## Abstract

**Background:** Galvanic vestibular stimulation (GVS) uses at least one electrode placed on the mastoid process with one or multiple placed over other head areas to stimulate the vestibular system. The exact electrode size used is not given much importance in the literature and has not been reported in several studies. In a previous study, we compared the clinical effects of using different electrode sizes (3 cm^2^ and 35 cm^2^) with placebo but with the same injected current, on postural control. We observed significant improvement using the smaller size electrode but not with the bigger size electrode. The goal of this study was to simulate the current flow patterns with the intent to shed light and potentially explain the experimental outcome.

**Methods:** We used an ultra-high-resolution structural dataset and developed a model to simulate the application of different electrode sizes. We considered current flow in the brain and in the vestibular labyrinth.

**Results:** Our simulation results verified the focality increase using smaller electrodes that we postulated as the main reason for our clinical effect. The use of smaller size electrodes in combination with the montage employed also result in higher induced electric field (E-field) in the brain.

**Conclusions:** Electrode size and related current density is a critical parameter to characterize any GVS administration as the choice impacts the induced E-field. It is evident that the higher induced E-field likely contributed to the clinical outcome reported in our prior study.

## Introduction

Galvanic Vestibular Stimulation (GVS) is a non-invasive stimulation technique that uses weak electrical current (< 5 mA) to modulate underlying vestibular afferent nerve fibers [1]. This technique has been used in various medical and non-medical applications for more than 100 years [1]. In medical research settings, GVS has shown promising results in diagnosing vestibular disorders as well as treating a wide range of nervous system diseases (e.g.,vestibulopathy, Meniere’s disease, gait abnormalities, etc.) [2]. GVS is characterized by different montages and by a constellation of parameters such a waveform type (e.g., DC, sine, sum of sines, noisy GVS), frequency, stimulation duration, etc [3–5]. Classical GVS produces stereotyped automatic perceptual postural and ocular responses by activating both primary otolithic neurons and primary semicircular canal neurons [1,2,6,7].

To characterize the safety and efficacy of any electrical stimulation technique, current density (i.e. current / area) is considered a critical technical parameter. High current densities are often associated with increased sensitivity but when threshold limits are exceeded, it can lead to brain injury [8]. While this is true, our group has also shown that appropriate choice of electrode material and design can be used to deliver high *electrode* current density without pain [9]. Herein, lies an important consideration that often gets overlooked and receives sparse attention in the literature. Electrode current density and the resulting *brain* current density, even though correlated due to the electrode placement, are intrinsically in-dependent stimulation metrics. In the context of a similar non-invasive electrical stimulation technique called transcranial direct current stimulation (tDCS), Nitsche and colleagues demonstrated that a current density value of 1 mA / 35 cm^2^ “standardizes” cortical response [10]. This aforementioned value, is however, closely related to the considered electrode montage and does not hold under all conditions. We discuss the potential limitation of this standardized current density value in Datta et al. [11]. Briefly, when two electrodes with the same injected electrode current density are placed immediately adjacent to each other (for instance, 1 cm away), dominant current flow shunts across the scalp leading to no underlying brain flow, thereby leading to no clinical / behavioral response. Another example is current delivery through electrophysiological electrodes (microelectrodes) matched to the standardized current density (0.0001 mA / 0.00035 cm^2^ same as 1mA / 35 cm^2^) will continue to be ineffective because of the inherently low current intensity value. Therefore, when comparing electrode montages with respect to corresponding effective dose delivery, it is worthwhile to match the brain (and not electrode) electric field (E-field) / current density.

Our group investigated the direct effect of electrode current density in a clinical study to shed more light on this issue [12]. For the first time, we considered two different electrode sizes in a GVS study. Specifically, we tested postural control in a 3 group parallel design (sham, subjects stimulated with a 3 cm^2^ electrode area and subjects stimulated with 35 cm^2^ electrode area). Each patient received 1 mA and therefore different current density was delivered to the two active groups. The stimulation duration was set to 30 minutes. For the sham group, current was ramped up to 1 mA for 30 seconds and then gradually decreased to mimic the skin sensation of the active arms. Our results indicated that postural stability can be significantly improved with the 3 cm^2^ size electrodes, but not with the 35 cm^2^ size electrodes. We postulated that the use of smaller electrodes perhaps resulted in more focal current delivery to the vestibular organs to explain the clinical outcome. The goal of the current study was to simulate current flow patterns under both of the above active conditions and to explore whether the use of smaller electrodes did indeed lead to more focal delivery. We consider an ultra-high-resolution model and determine current flow pattern on the cortical surface, and the two implicated structures for GVS mechanism of action (otolith and the semicircular canals).

## Methods

The ultra-high resolution head (MIDA: Multimodal Imaging-Based Detailed Anatomical Model) available through the IT’IS Foundation was used in this study [13]. This dataset was suitable for this study because its resolution (500 μm isotropic) allows visualization and therefore analysis of the small vestibular regions of the head. The simulation workflow was largely based on previous work of our group [14–16]. These steps were:

1. Image-processing and segmentation of the data. First, the nifty (.nii) color masks from the MIDA dataset were processed in MATLAB to generate corresponding tissue masks based on the intensity values. The resulting masks were imported into Simpleware (Synopsys Ltd., CA, USA) to correct anatomical and continuity errors. Masks containing similar electrical conductivity (e.g. the skull dipole, outer table, and the mandible make up the bone mask) were combined to form a single mask, except for the areas of interest (e.g., vestibular area) through which current flow analysis was to be performed.
2. Replicating electrode geometry and placement. Two different electrode sizes were modeled in Simpleware to mimic the 3.14 cm^2^ (1 cm radius) and 35 cm^2^ (5 × 7 cm) application used experimentally (Fig 1). The geometry of the electrodes were circular and rectangular, respectively. We refer to the smaller electrode as a 3 cm^2^ electrode for simplicity - similar to the clinical study [12]. The shape of the gel remained the same as that of the electrodes. The thickness ratio for both electrode sizes relative to the gel was 0.5:1. The electrodes and gel were then placed on the mastoid area of the skin tissue mask in a Bilateral-Bipolar configuration as illustrated in Fig 1.
3. Meshing and Finite element method (FEM) model generation. The two different models (corresponding to the two electrode sizes) and each of the segmented tissue compartments/masks were adaptively meshed using Simpleware. The mesh was imported into COMSOL Multiphysics 5.6 (COMSOL Inc., MA, USA) to set up a FEM model to compute current flow and to subsequently generate electrical field (E-field) surface plots of the regions of interest. The isotropic and homogeneous electrical conductivity value in S/m assigned to each mask were: skin (0.465), skull (0.01), cerebrospinal fluid (CSF) (1.65), gray matter (0.276), white matter (0.126), air (1e-7), cranial nerves (0.017126), ear auricular cartilage (0.16113), ear semicircular canals (2), blood (0.7), gel (1.4), and electrodes (5.8e7). The noisy GVS waveform considered in our clinical study involved a spectrum ranging from 0 to 640 Hz [17]. Simulating application of any waveform with frequency content requires consideration of a modified Laplace equation incorporating a reactive component [18]. However, the real component (i.e. conductivity) dominates at frequencies up to 10 kHz, implying that the standard Laplace equation can still be used. The model physics was therefore formulated with standard Laplace with the following boundary conditions: 1) normal current density condition for the left (anode) electrode corresponding to 1 mA, 2) ground at the right (cathode) electrode, and 3) all external surfaces treated as insulated.

**Figure 1.**
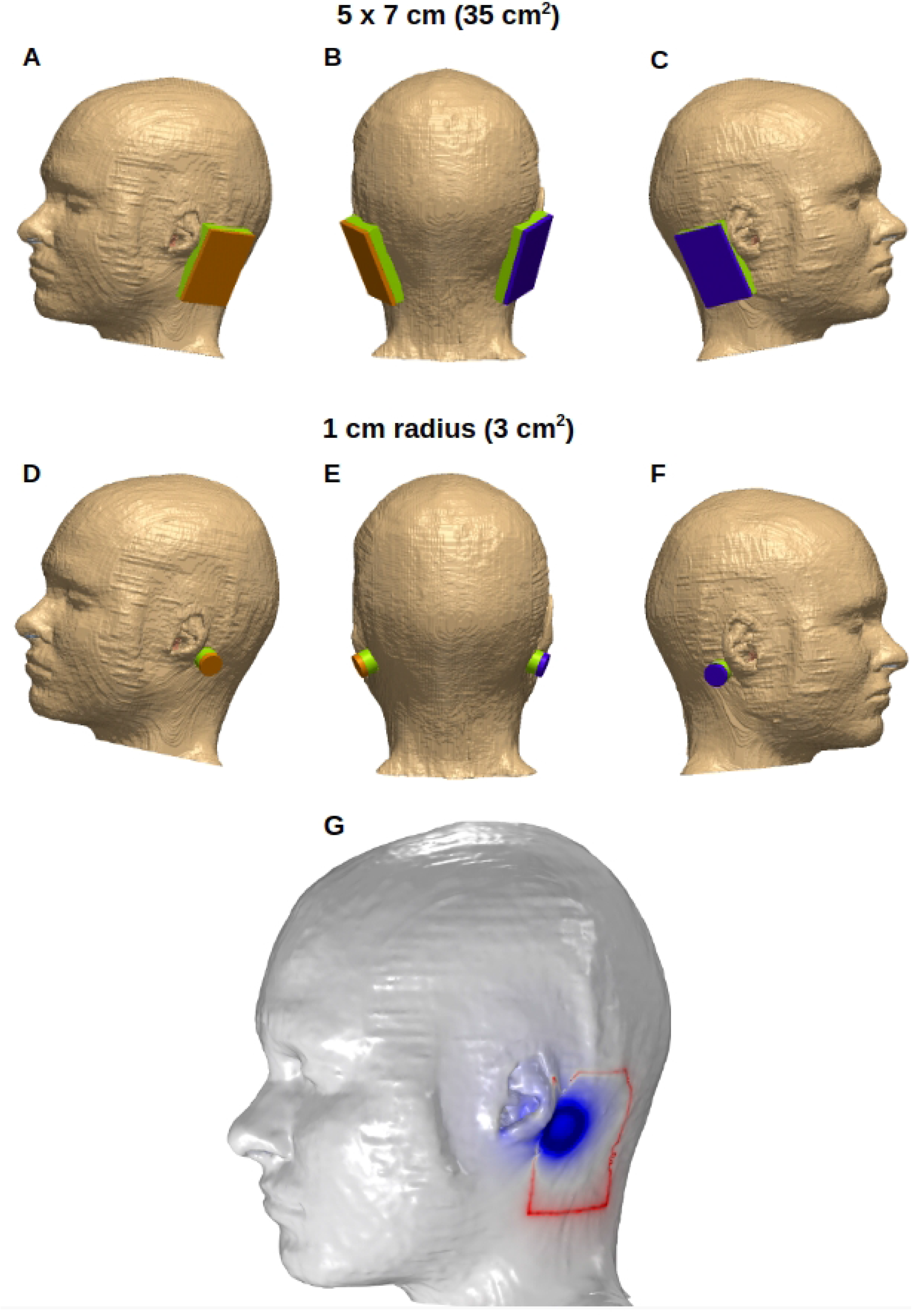
Electrode Placements. Two different electrode sizes were simulated to apply GVS: 35 cm^2^ (A-C) and ~3 cm^2^ (D-F). The electrodes were placed on the skin-tissue mask in a Bilateral-Bipolar configuration. (G): The two electrode sizes were overlaid on the same head model geometry to highlight final placement with respect to the anatomy.

## Results

For each model considered in this study, the induced E-field / current density plots of the brain and vestibular regions were obtained, as shown in Fig 2. We further overlaid current flow streamlines induced due to both models (Fig 2-C1) and generated a difference plot on the regions of interest (Figs 2-C2 and 2-C3) to analyze potential differences in a systematic fashion. The current flow profile of the Bilateral-Bipolar montage is lateral (left to right) as expected. The simulations confirm that the current entering from one mastoid location at the anode electrode traverses all intermediate tissues until it reaches the brain, and then exits at the second mastoid location at the contralateral cathode electrode side. Figs. 2-A2 and 2-B2 indicate that the dominant current flow occurs at the cerebellum and the brain stem region, with some flow in the temporal regions for the model using smaller electrodes. Further, current flow is largely minimal (< 0.1 V/m) in the frontal, parietal and motor cortical regions of the brain.

**Figure 2.**
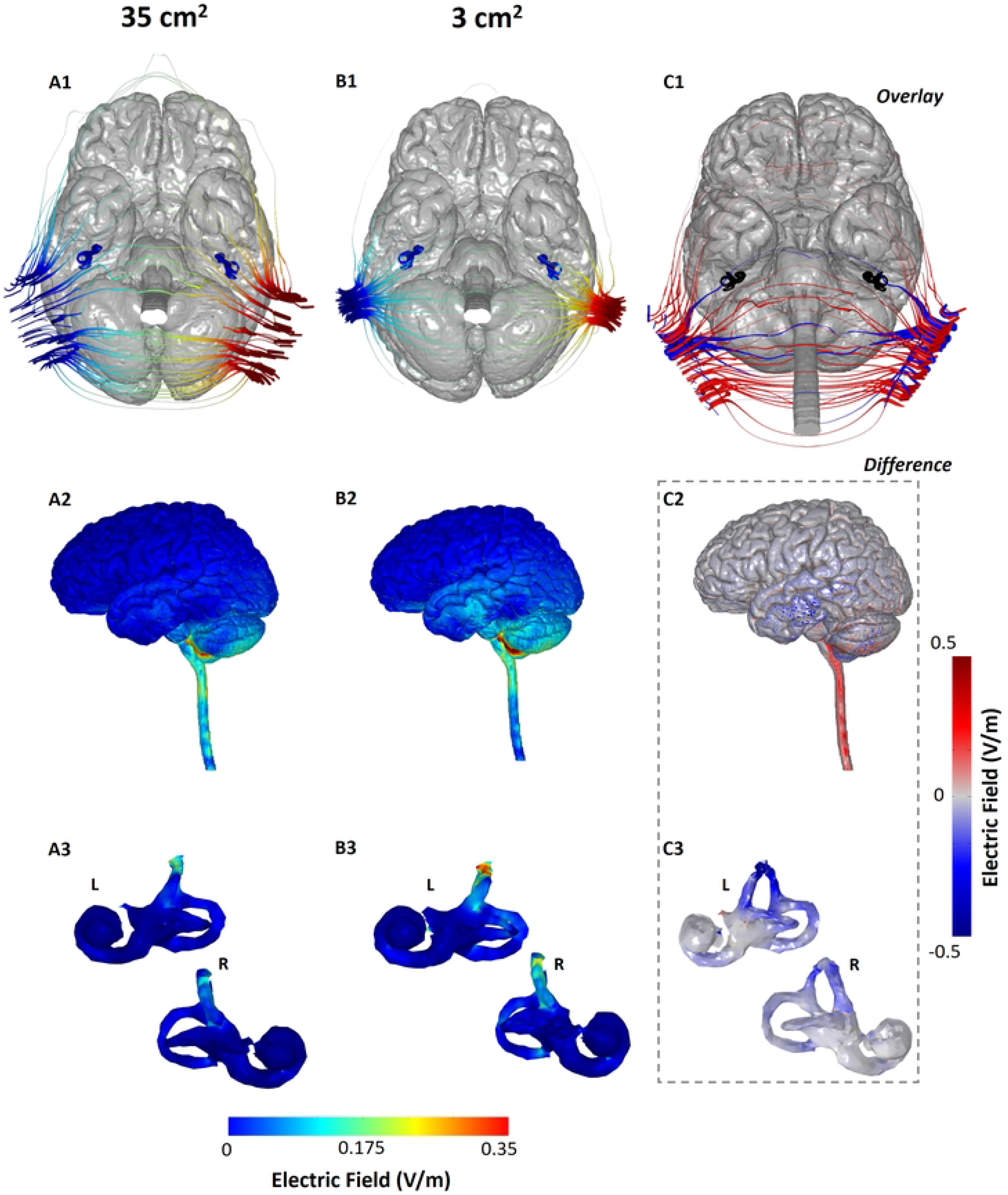
Comparison of GVS induced current flow using small- and large-size electrodes. Columns A (A1, A2, A3) and B (B1, B2, B3) correspond to the 35 cm^2^ and 3 cm^2^ electrodes, respectively. The current streamlines from both models are overlaid in C1. C2 and C3 indicate the difference in induced E-field due to the two electrode sizes. The first row (A1, B1 and C1) shows the current streamline plots with the brain, SCC and otolith organs masks visible to provide enhanced visualization of current flow with respect to their anatomy. The second row (A2, B2, and C2) shows the induced brain surface E-field plots. Similarly, the third row (A3, B3, and C3) shows the induced E-field plots on the vestibular labyrinth.

The predicted maximum E-field (based on the 99^th^ percentile) delivered to the brain using 3 cm^2^ electrodes is 0.175 V/m (Fig 2-B2) and the maximum E-field delivered to the brain using 35 cm^2^ electrodes is 0.147 V/m (Fig 2-A2). The current streamlines help visualize the entry of current through the mastoid region using both models (Figs. 2-A1 and 2-B1). It can be clearly seen that in the case of the 3 cm^2^ electrodes, a restricted entry point leads to an increased concentration of the lines of force at the mastoid locations. Additionally, smaller-sized electrodes result in a greater separation between the electrodes (due to greater edge to edge or perimeter to perimeter distance), which results in the current having a lower “motivation’’ to flow around the back or around the front of the head. Therefore, the net effect of delivering current through smaller-sized electrodes is not only focal delivery, but higher current intensity reaching regions of interest in the brain. In the case of the 35 cm^2^ electrodes, the current entry in the mastoids by default is not as concentrated and, due to the smaller effective distance between the electrodes, there is more “current loss” from the back of the head than from the front of the head. This phenomena is clearly highlighted by the higher number of streamlines running from one electrode to the other across the back of the head (Fig 2-A1) and some running around the front of the head. As well, this is further emphasized by the overlay images of the two simulations (Fig 2-C1) that illustrates the focal entry and dominant lateral flow of the blue (3 cm^2^) as opposed to the red streamlines (35 cm^2^).

While current input is restricted at the level of the skin and the skull, we actually observe higher current flow in the temporal cortex using the 3 cm^2^ electrodes. This is counter-intuitive but supported by the overall current flow pattern due to any transcranial electrical stimulation modality. The scalp and the CSF are the most conductive tissues in the head. When current is first injected at the level of the scalp, there is some shunting (i.e. flow *across* the skin rather than *into* the tissue) due to its higher conductivity with respect to the underlying skull [9]. The extent of this shunting is determined by the electrode montage. The remaining current entering the skull largely flows into the next tissue (CSF), as there is less incentive for current to flow across the low conductive skull. Since less current shunt (loss) occurs at the scalp level in the 3 cm^2^ electrode, the magnitude of the current entering the CSF is higher compared to the current entering the CSF with the 35 cm^2^ electrodes. In summary, a combination of the aforementioned factors explains why higher current flow is observed at the temporal cortex due to the smaller sized electrodes. For the larger electrodes, as there is more current loss across the scalp, there is more inferior current flow. This is evident in the observed E-field magnitude (~0.3 V/m) in regions extending along the spinal cord. This is further emphasized via the difference image (Fig 2-C2) that indicates that the 3 cm^2^ electrode model results in more flow in the cerebellum and the temporal regions (blue hue), whereas, the 35 cm^2^ electrode model results in more flow in the brain stem regions (red hue).

When evaluating flow at the level of the vestibular end organs, similar flow (spatial distribution) but higher induced E-field (Fig 2-B3) was observed for the model simulating smaller electrodes. For both models, we observe increased current flow in the anterior canals with respect to the lateral and posterior canals. The maximum E-field delivered to the left vestibular network with the 35 cm^2^ and 3 cm^2^ electrodes was 0.126 V/m and 0.201 V/m respectively. Similarly, when considering the right network, the induced E-field is 0.108 V/m when using 35 cm^2^ electrodes and 0.165 V/m when using the 3 cm^2^ electrodes. The relative E-field percentage increase (Table 1) indicates a slightly higher value for the left network. This can be expected as the head anatomy is not perfectly symmetric (with respect to the sagittal plane) and therefore, variations in current flow will manifest in different percentage increases.

**Table 1.** Maximum induced E-field (based on 99th percentile) in the brain and vestibular regions across the two models. The percentage increase of E-field with respect to the larger electrode model is noted in the final column.

| Electric field (V/m) | 35 cm<sup>2</sup> | 3 cm<sup>2</sup> | (E<sub>3</sub>-E<sub>35</sub>) / E<sub>35</sub> |
| --- | --- | --- | --- |
| Brain (White and gray matter) | 0.147 | 0.175 | 19.05% |
| Left vestibular region | 0.126 | 0.201 | 59.52% |
| Right vestibular region | 0.108 | 0.165 | 52.78% |

## Discussion

The predictions in this study verify the focality increase that we previously proposed as the primary reason for the postural control improvement using smaller size electrodes [12]. The concentrated current entry into the head in combination with the montage employed, in fact, results in more current entering the brain — resulting in higher induced E-field in the brain and the vestibular labyrinth. While induced E-field magnitude in the target of interest is one of the main drivers of neuromodulation, a one-to-one linear translation of clinical / behavioral effects is not expected due to several factors. It is for this reason that clinical effects with 2 mA for instance, are not expected to simply double from the effects observed at 1 mA - as observed from tDCS studies [19]. Nonetheless, it is well accepted that neuronal modulation is directly related to induced E-field. This has been extensively demonstrated by a multitude of experimental and theoretical studies indicating that polarization of short or bent cortical axons (20-21) and synaptic efficacy vary linearly with E-field magnitude (22) and neuronal excitability metrics vary with E-field magnitude (22,23). We are therefore able to conclude that the higher induced E-field contributed in some fashion to the improvement in postural stability. We also note that the increases in induced E-field in cortical (19%) and vestibular regions (52- 59%) with the smaller electrodes are in the same order of percentage improvement of the clinical effects. While a direct correspondence cannot be expected, our results offer a plausible explanation of the experimental findings in Nooristani et al. [12]. In summary, we demonstrate the importance of electrode size and encourage the community to document dimensions in their publications.

Our finding that the anterior canals receive greater current flow with the Bilateral-Bipolar montage (irrespective of the electrode size used) would imply greater sensitivity of the canal with respect to the lateral and the posterior canal. This finding could potentially be used to further adjust weights of the vector summation model proposed by Day et al. [24]. In general, a future research topic could be to use current flow model predictions as used here across different GVS electrode montages to test resultant impact on the vector summation model’s accuracy. We also note that calibration of current intensity is frequently used in GVS administration where individualized intensity is delivered based on the subjects’ perceptual threshold. The static field approximation in our model implies linearity of the solution and therefore induced E-field will scale linearly with the stimulation magnitude. So the induced cortical E-field for 0.7 mA scalp current will be simply 0.7 times the cortical E-field induced for 1 mA. In summary, computational modeling can be used to support GVS administration in several ways similar to efforts transforming other modalities (DBS, TMS, etc) [25,26]. As research using GVS increases and evolves, there is potential to leverage modeling to help with determining optimal parameters (electrode location, electrode shape, etc.), relate current flow to stimulation outcome, perform safety analysis, etc. We expect this study to help demonstrate this utility and encourage additional efforts.

## Author Contributions

DQT and AD developed the concept idea. DQT and AG performed the E-field modeling and related post-processing. DQT, AD, MN, MM, and FC confirmed the overall methodology. All authors contributed to writing the article and approved the submitted version.

## Funding

The work performed in this study was partially funded by grants to AD from DoD-DARPA (W912CG21C0014)

## Conflicts of Interest

DQT, AG, and AD are employees of Soterix Medical. The remaining authors declare that the research was conducted in the absence of any commercial or financial relationships that could be construed as a potential conflict of interest.

*-

